# VAPiD: a lightweight cross platform viral annotation pipeline and identification tool to facilitate virus genome submissions to NCBI GenBank

**DOI:** 10.1101/420463

**Authors:** Ryan C. Shean, Negar Makhsous, Graham D. Stoddard, Michelle J. Lin, Alexander L. Greninger

## Abstract

**Background:** With sequencing technologies becoming cheaper and easier to use, more groups are able to obtain whole genome sequences of viruses of public health and scientific importance. Submission of genomic data to NCBI GenBank is a requirement prior to publication and plays a critical role in making scientific data publicly available.

GenBank currently has automatic prokaryotic and eukaryotic genome annotation pipelines but has no viral annotation pipeline beyond influenza virus. Annotation and submission of viral genome sequence is a non-trivial task, especially for groups that do not routinely interact with GenBank for data submissions.

**Results:** We present Viral Annotation Pipeline and iDentification (VAPiD), a portable and lightweight command-line tool for annotation and GenBank deposition of viral genomes. VAPiD supports annotation of nearly all unsegmented viral genomes. The pipeline has been validated on human immunodeficiency virus, human parainfluenza virus 1-4, human metapneumovirus, human coronaviruses (229E/OC43/NL63/HKU1/SARS/MERS), human enteroviruses/rhinoviruses, measles virus, mumps virus, Hepatitis A-E Virus, Chikungunya virus, dengue virus, and West Nile virus, as well the human polyomaviruses BK/JC/MCV, human adenoviruses, and human papillomaviruses. The program can handle individual or batch submissions of different viruses to GenBank and correctly annotates multiple viruses, including those that contain ribosomal slippage or RNA editing without prior knowledge of the virus to be annotated. VAPiD is programmed in Python and is compatible with Windows, Linux, and Mac OS systems.

**Conclusions:** We have created a portable, lightweight, user-friendly, internet-enabled, open-source, command-line genome annotation and submission package to facilitate virus genome submissions to NCBI GenBank. Instructions for downloading and installing VAPiD can be found at https://github.com/rcs333/VAPiD.

## Background

With sequencing technologies becoming cheaper and more accessible, genomic sequencing data is becoming increasingly widespread. Smaller groups are generating more sequencing data than they can analyze alone. In order to extract maximal scientific and public health value out of these data, sharing of assembled consensus genomes and raw sequence data is critical. The democratization of genomics takes a village. This is especially true for infectious diseases, where the searchable sequence databases allow for real-time tracking of viral epidemics and solving of foodborne bacteria outbreaks [1–3]. While recent high-profile cases receive the most attention, the fact remains that almost all infectious diseases exist in the context of an ongoing outbreak [1,4]. In the clinical world, metagenomic analysis pipelines depend on and take advantage of the availability of many infectious disease genomes to allow for faster and accurate alignments [5–7]. In the basic science world, a graduate student studying the function of a singular protein is greatly assisted by being able to pull the world’s history of sequence diversity for that protein before designing experiments [8].

Many foresee a world in which nearly every infectious disease genome is sequenced and archived in a publicly searchable database [9,10]. State and federal public health laboratories have built capacity such that they now sequence more than 6,000 influenza virus genomes and more than 5,000 enteropathogenic bacterial genomes each year [11]. Major efforts in rationalizing workflows, from nucleic acid extraction to data deposition and analysis, have enabled these rapid growths in throughput [12–14]. These tools allow public health and research laboratories to focus on the epidemiological or scientific insight gained from sequencing rather than rote protocols for data deposition. Specifically in the area of genome annotation, the National Center for Biotechnology Information (NCBI) has created the Prokaryotic Genome Annotation Pipeline, Eukaryotic Genome Annotation Pipeline, and the Influenza Virus Sequence Annotation Tool [15,16].

Surprisingly, other than for influenza virus, NCBI GenBank does not currently have an automatic viral genome annotation pipeline. The incredible diversity of DNA and RNA viruses presents a challenge for development of a universal annotator [17]. Complex viral life cycles involving RNA editing, ribosomal slippage, and overlapping reading frames create additional annotation issues, along with non-standard nomenclature for viral gene products [18–20].

In order to accept submitted viral genomic data, NCBI GenBank requires 1) viral sequence complete with at least one protein annotation, 2) author/depositor metadata, and 3) viral sequence metadata, such as strain, collection date, collection location, and coverage. While manual annotation of nucleotide sequence can be done for small numbers of viruses, it is extremely time-consuming and labor-intensive. Even after correct annotations have been obtained for all viral sequences, manually integrating author and sample metadata to create files to be submitted is an equally time-consuming effort and not a feasible solution for groups sequencing more than a few viruses.

To date, existing viral annotation tools have focused largely on batch submissions of a single virus species. This may be a holdover from when specific PCR-based methods were required to faithfully recover viral whole genome sequence or due to the focus of researchers on a single virus at a time. With the increasing use of metagenomic or shotgun next-generation sequencing and the availability of more and more sequencing capacity, researchers can confidently batch many different RNA or DNA viruses together on a single sequencing run.

In order to facilitate viral genome annotation, we have developed a lightweight and user-friendly command-line tool that inputs FASTA files of complete or near complete viral genomes, automatically annotates them, and outputs the required files for GenBank submission over email. VAPiD handles batch submissions of multiple viruses of different types without prior knowledge of the viral species, correctly annotates RNA editing and ribosomal slippage, performs spellchecking on annotations, handles batch or individual submission of metadata, runs with a simple one-line command, and creates annotated viral sequence files for GenBank submission.

### Implementation

VAPiD can be downloaded at https://github.com/rcs333/VAPiD. An installation guide, usage instructions, and test data can also be found at the above website. The invocation of VAPiD is shown in Figure 1, users must provide a standard FASTA format containing all of the viral genomes they wish to annotate. Users also must provide a GenBank submission template (.sbt file) that includes author, publication, and project metadata. The GenBank submission template can be used for multiple viral sequences or submissions and is easily created at the NCBI Submission portal (https://submit.ncbi.nlm.nih.gov/genbank/template/submission/). An optional sample metadata file (.csv file) can also be provided to VAPiD to expedite the process of incorporating sample metadata. This optional file can also be used to include any of the Source Modifiers supported by NCBI (https://www.ncbi.nlm.nih.gov/Sequin/modifiers.html). If no sample metadata file is provided, VAPiD will prompt the user to input the required sample metadata at runtime. Additionally, users can provide a specified reference from which to annotate all viruses in a run, as well as provide their own BLASTn database or force VAPiD to search NCBI’s NT database over the internet.

**Figure 1.**
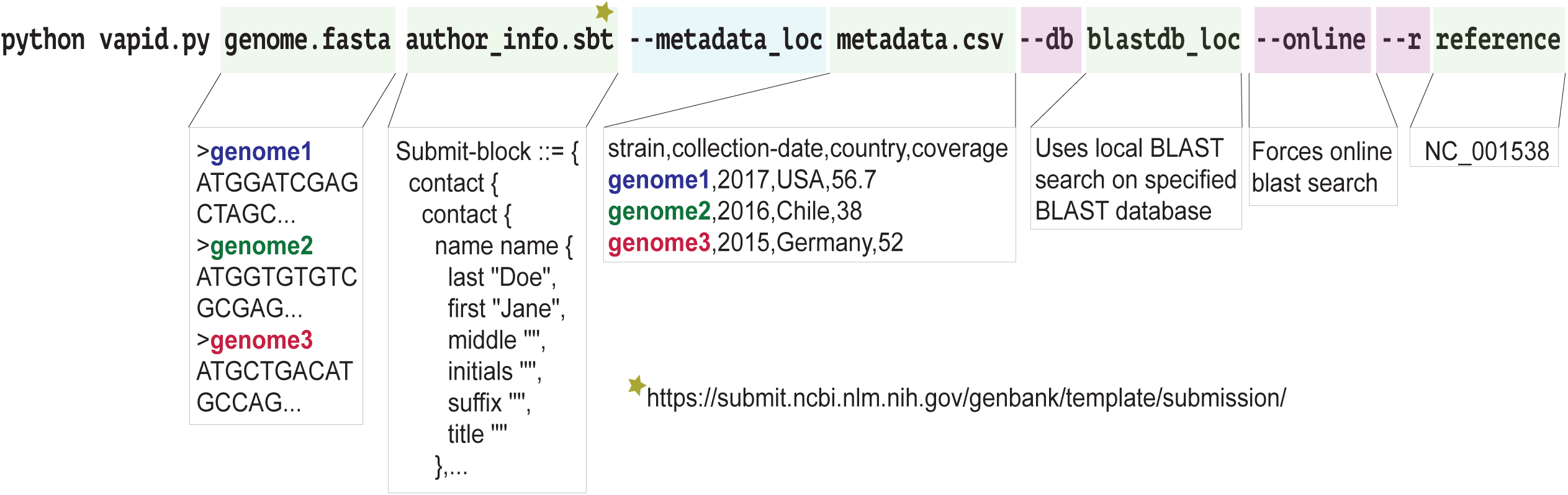
Example usage of VAPiD. The two required files are shown as genome.fasta and author_info.sbt. genome.fasta is all of the viral genomes you wish to submit, named as you want them to appear on GenBank. In the example code provided in the github repository this example file is called example.fasta. The author_info.sbt file is an NCBI specific file for attaching sequence author names to sequin files and is a required part of properly submitting sequences to NCBI. This file can be generated at (https://submit.ncbi.nlm.nih.gov/genbank/template/submission/). The first optional command is a comma separated file in which you can include all relevant metadata. You can create additional columns here so long as they correspond to NCBI approved sequence metadata. A list and formatting requirements can be found at (https://www.ncbi.nlm.nih.gov/Sequin/modifiers.html). Note that FASTA sequence names must be identical to names in the optional metadata sheet. Additionally, one could omit the metadata sheet and VAPiD will prompt for strain name, collection-date, country, and coverage data automatically at runtime. The second optional argument is a location of a local BLASTn database, which will force VAPiD to use the specified database instead of the included database. The last optional argument will force VAPiD to send an online search query to NCBI’s NT database.

The VAPiD pipeline is summarized in Figure 2. The first step is finding the correct reference sequence. This is accomplished in three ways 1) using the provided reference database (default), 2) forcing VAPiD to execute an online BLASTn search of NCBI’s NT database, or 3) inputting the accession number of a single NCBI sequence to use as the reference.

**Figure 2.**
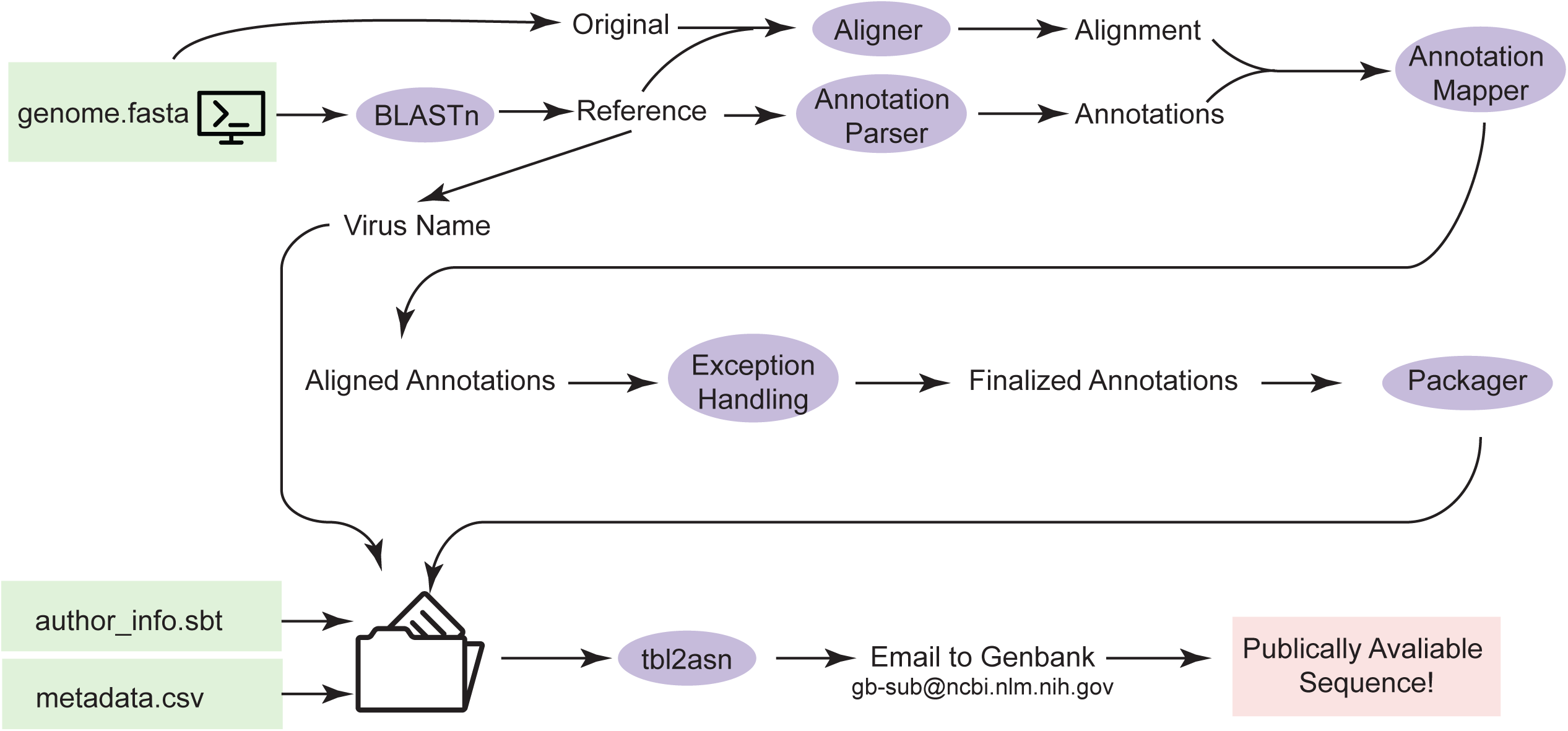
General design and information flow of VAPiD. First the provided sequences are used as queries for a local BLAST search (default) or an online BLASTn search. After results have been returned a reference annotation is downloaded, if a specific reference accession number is given than this reference is downloaded. Then the original FASTA file is aligned with the reference FASTA and the resulting alignment is used to map the reference annotations onto the new FASTA. Then custom code runs through the file and handles RNA editing, ribosomal slippage and splicing. These finalized annotations are then plugged into NCBI’s tbl2asn with the author information and sequin files are generated as well as gbk files which can be used to manually verify accuracy of new annotations. Quality checked .sqn files can be emailed directly to GenBank.

In the default case, NCBI’s BLAST+ tools are called from the command line to search against a reference database that is included with the VAPiD installation. This database was generated by downloading all complete viral genomes in NCBI on May 1, 2018. The best result from this search is passed as the reference into the next steps.

If the online option for finding the reference is specified, VAPiD finds an appropriate reference sequence for each genome to be annotated by performing an online BLASTn search with a word size of 28 using BioPython’s NCBIWWW.qblast() function against the online NCBI NT database. The BLASTn output is parsed for the best scoring alignment among the top 15 results that contains “complete genome” in the reference definition line. If no complete genome is found in the top 15 BLASTn results, it uses the top-scoring hit as the reference sequence.

If a specific reference is provided, VAPiD simply downloads it directly from NCBI. After the correct reference is downloaded, gene locations are stripped from the reference and a pairwise nucleotide alignment between the reference and the submitted sequence is generated using MAFFT [21]. The relative locations of the genes on the reference sequence are then mapped onto the new sequence based off the alignment. This putative alignment only requires that start codons are in regions of high homology and does not rely on intergenic spacing or gene lengths. Gene names are taken from the annotated reference sequence GenBank entry. Automatic spellchecking is performed with NCBI’s ESpell module. This module provides automatic spellchecking of many biological strings including protein product names. An optional argument can be provided at execution that disables this automatic step.

The diverse array of methods viruses use to encode genes can present problems for any viral genome annotator. Ribosomal slippage allows viruses to produce two proteins from a single mRNA transcript by having the ribosome ‘slip’ one or two nucleotides along the mRNA transcript, thus changing the reading frame. Since ribosomal slippage is well conserved within viral species and complete reference genomes often list exactly where it occurs, custom code was used to strip the correct junction site and include it in the annotation.

RNA editing is another process by which viruses can include multiple proteins in a single gene. In RNA editing, the RNA polymerase co-transcriptionally adds one or two nucleotides that are not on the template. These changes are specifically created during viral mRNA transcription and not during viral genome replication. RNA editing presents an annotation issue because the annotated protein sequence does not match expected translated nucleotide sequence. To correctly annotate genes with RNA editing, VAPiD parses the reference genome viral species, detects the RNA editing locus, and mimics the RNA polymerase. VAPiD adds the correct number of non-templated nucleotides for the viral species and provides an alternative protein translation. This process is hard-coded for human parainfluenza 2-4, Nipah virus, Sendai virus, measles virus, and mumps virus. Although RNA editing occurs in Ebola virus, references for Ebola virus are annotated in the same way as ribosomal slippage, so code written for ribosomal slippage handles Ebola virus annotations.

After ribosomal slippage and RNA editing are processed, files required for GenBank submission are generated with the provided author and sample metadata. VAPiD first generates the .fsa file, .tbl file, and optional .cmt file. Submission files for each viral genome are packaged into a separate folder for each sequence. VAPiD then runs tbl2asn on each folder using the provided GenBank submission template file (.sbt). tbl2asn generates error reports and Sequin (.sqn) and GenBank (.gbk) files for manual verification and GenBank submission via email attachment to.

## Results and Discussion

To illustrate our vision of how VAPiD will be useful to the scientific community we are providing two example use cases.

### Case Study 1

This first example is the task that the authors originally wrote VAPiD for - annotating large numbers of genomes from different viral species, which mirrors the type of data that many clinical and public health laboratories may encounter. To illustrate this, 30 synthetic viral genomes were created by manually mutagenizing sequences downloaded from NCBI GenBank. Only point mutations that did not lead to stop codons were used and each sequence was given about 10 - 25 changes depending on the viral genome length, equal to roughly one amino acid change per 1,000 nucleotides. The species included were Nipah, Sendai, Measles, Mumps, Parainfluenza 1-4, Ebola, Rotavirus segments, MERS, SARS, Coronavirus 229E, West Nile Virus, HTLV, HIV1, Hepatitis A-C, Hepatitis E, Norovirus, Enterovirus, JC virus, and BK polyomavirus. All genomes were put together into a single FASTA file. Metadata (collection date, country and coverage) for each of the viral genomes was put together into a single csv file. VAPiD was then executed with the command [python vapid.py example.fasta example.sbt --metadata_loc example_metadata.csv]. After running for 123 seconds the program successfully generated 30 complete and correct NCBI submittable annotation files (*.sqn), which could be immediately emailed to NCBI. When these same sequences were run with the online option it took 280 minutes to complete, most of which was waiting for NCBI to return BLAST results. The above example shows how VAPiD would be ideal for busy groups who produce a diverse array of viral sequence such as public health or clinical testing labs.

### Case Study 2

The second example use case highlights both the cross platform functionality of VAPiD as well as the ability to manually enter NCBI required metadata at run time and use an online BLAST search - reducing the number of required files at runtime to two. For this example, a sample human parainfluenza 3 virus FASTA was created. No accompanying metadata file was created (unlike in the first example). Additionally, the syntax is exactly the same to use VAPiD across all operating systems. The single human parainfluenza 3 virus took approximately two minutes to annotate (including waiting about 30 seconds for NCBI blast results to be returned) on a Windows 10 Virtual machine with a single 2.36 GHz core and 2 Gb of RAM. This example highlights how VAPiD can be used to annotate and submit viruses with only two input files (the author template file only needs to be created once) on almost any Mac, Linux, or Windows computer with a python installation.

### Comparisons

Two other programs exist to handle viral annotation, JCVI’s VIGOR (https://doi.org/10.1186/1471-2105-11-451) and the Broad Institute’s viral-ngs suite (https://viral-ngs.readthedocs.io/en/latest/) (Table). VIGOR is an extremely fast and automatic viral annotator that is able to correctly annotate any number of virus genomes without prior knowledge of viral species or type. VIGOR also is able to correctly handle RNA editing and ribosomal slippage – needing only an input FASTA file with the genome to be annotated, it can produce NCBI compatible .tbl annotation files. VIGOR works natively on Mac and Linux systems. The annotations that VIGOR produces are identical to those created by VAPiD except that VAPiD only annotated CDS regions whereas VIGOR will annotate all regions. Two primary advantages VAPiD offers over VIGOR are 1) VAPiD can be installed and run on Windows systems, and 2) VAPiD automates metadata and NCBI submission preparation steps. This fits our intended use case of a public health laboratory that may only own Windows machines and does not want to spend extra time worrying about scripting tbl2asn and manually preparing metadata files. For those with powerful Linux machines and some knowledge of bash scripting we recommend stitching VIGOR and tbl2asn together with custom scripts. However, for those without the infrastructure or time to pursue this path VAPiD offers a reasonable alternative.

The Broad Institute’s publicly available viral-ngs package, also developed in Python, represents another alternative to VAPiD. This suite of tools is custom developed for internal Broad Institute users but publicly available and well-documented on GitHub. Viral-ngs takes a reference genome and corresponding annotation file and transfers the reference annotations onto a new genome. This is very similar to the way VAPiD transfers annotation, except viral-ngs requires the user to provide a reference, which avoids the potential problem of low quality references propagating errors. Viral-ngs is well suited for annotating large numbers of the same type of virus all at the same time. Another advantage of viral-ngs is that it contains many tools for going from raw sequencing reads to complete viral genomes. While viral-ngs handles ribosomal slippage and most normal viral genes, because NCBI compatible annotation files do not contain enough information to correctly annotate RNA editing viral-ngs fails in these cases. Outside of cases involving RNA editing VAPiD and viral-ngs produce identical CDS annotations, with viral-ngs transferring all other features. This software, like VAPiD, is also cross platform and could be easily run on a Windows machine with a slightly modified Python installation. The main advantages VAPiD offers over viral-ngs are the ease of batching multiple viral types together and also as with VIGOR, the automated metadata input and NCBI packaging.

### Limitations

As with all software tools and especially gene annotation programs, VAPiD is not without limitations. VAPiD was designed purely to expedite the process of submitting large batches of different human virus genomes from a clinical laboratory. As such, a major limitation is that VAPiD expects a “complete genome” to use as a reference to be available for each of the viral sequences submitted to it. VAPiD is not the preferred annotation tool for novel or extremely divergent viral species. However, we find that complete reference genomes are available for most viral species of clinical importance, especially for viruses that are commonly sequenced in clinical or public health laboratories. A further limitation of VAPiD is that it does not perform *ab initio* gene annotation. Instead VAPiD simply transfers annotations from the closest reference genome with a few quality control steps, such as allowing for slightly different gene lengths and RNA editing. A significant limitation of this strategy is that because the new annotation is generated from a downloaded reference, any errors that are in the downloaded reference will be transferred to the new genome. This means that VAPiD performs best on high-quality and accurate reference sequences. The downside of the VAPiD strategy is that if an inaccurate reference strain is used, error can propagate extremely fast. For example, early in our development we deposited roughly 60 human parainfluenza 3 viruses that had the matrix protein incorrectly annotated as ‘matrix potein’ [sic] due to a misspelling in the official NCBI reference sequence NC_001796 for HPIV3. To combat this problem, we have included a variety of ways to pull references, including directly specifying the reference. However, due to ease of implementation this functionality only works if all the viruses to be submitted are the same type. For those submitting many viruses of the same type we recommend using this feature to ensure annotation quality. VAPiD also contains code that overwrites misspellings in protein product names using NCBI’s ESpell utility. This spell-checking step automatically flags and corrects errors such as the misspelling mentioned above. A helpful message of what was corrected is printed to the console to prevent erroneous corrections. While this takes care of most misspellings, it is not a perfect fix, and viruses that we tested previously could have erroneous references uploaded in the future and further propagate error. It is for this reason that we highly recommend that users inspect and verify the accuracy of their new annotations, which have to potential to be used for future references.

## Conclusions

VAPiD allows users to go from any number of viral genome sequences to GenBank-ready submission files (.sqn) with a single command. VAPiD runs from the command line with two required arguments and one optional sample metadata file, with the option of inputting minimal sample metadata via the command line at the time of annotation. The latest version of VAPiD can be found at https://github.com/rcs333/VAPiD along with detailed installation and usage instructions. This software will allow public health, clinical virology, and research laboratories that do not have the resources to develop their own in-house genome annotation and submission tools or to adapt other pre-existing tools to quickly and easily share the sequence information that they generate on a regular basis with a minimal effort.

## Availability and Requirements

Project name: VAPiD

Project Homepage: https://github.com/rcs333/VAPiD

Operating System(s): Mac OS X, Linux, Windows 10

Programing Language: Python

Other requirements: Python 2.7.4 or higher, BioPython, tbl2asn, MAFFT, internet connection

License: MIT

Any restrictions to use by non-academics: None

## List of Abbreviations

NCBI: National Center for Biotechnology Information
BLAST: Basic Local Alignment Sequent Tool
VAPiD: Viral Annotation Pipeline and iDentification
sqn: Sequin file
sbt: Submission Template file
csv: Comma separated value file

## Declarations

### Ethics approval and consent to participate

Not applicable

### Consent for publication

Not applicable

### Availability of data and material

All data generated or analyzed during this study are included in this published article, GitHub, or NCBI BioProject PRJNA338014. Code and databases are available on GitHub (https://github.com/rcs333/VAPiD)

### Competing interests

The authors declare that they have no competing interests.

### Funding

This project was supported by the University of Washington Department of Laboratory Medicine and Virology Division. Ryan Shean was funded by the Mary Gates Research Scholarship at the University of Washington.

### Authors’ contributions

RS wrote the code. NM generated sequences for beta-testing and development of the software. ML helped implement several features to correctly annotate RNA editing and ribosomal slippage. GS implemented cross-platform functionality. AG conceived of the software and oversaw its development. RS and AG wrote the paper.

All authors contributed to debugging and testing of the final version of the software and all authors read and approved the final manuscript.

**Table** – Attribute comparison of VAPiD, VIGOR, and viral-ngs annotation pipelines

